# Real-world objects scaffold visual working memory for features: Increased neural engagement when colors are remembered as part of meaningful objects

**DOI:** 10.1101/2025.04.15.645785

**Authors:** Yong Hoon Chung, Timothy F. Brady, Viola S. Störmer

## Abstract

Visual working memory is a core cognitive function that allows active storage of task-relevant visual information. Contrary to the common assumption that the capacity of this system is fixed with respect to a single feature dimension, recent research has shown that working memory performance for a simple visual feature – color – is improved when this feature is encoded as part of a real-world object relative to an unrecognizable scrambled object. Using EEG (N = 24), we here demonstrate that this performance benefit is supported by increased neural engagement during the retention period, as indexed by enlarged contralateral-delay-activity during maintenance. Furthermore, the pattern of neural activity across parietal-occipital electrodes was more stable across time, suggesting that real-world objects may support more robust memory representations. Finally, we report a novel fronto-central event-related potential that distinguishes between real-world objects and scrambled objects during encoding and maintenance processes. Overall, our results demonstrate that active visual working memory capacity for simple features is not fixed but can expand depending on what context these features are encoded in.

## Introduction

Visual working memory supports the active storage of task-relevant visual information (Baddeley & Hitch, 1974; Bays et al. 2024). Its capacity is typically assessed by presenting multiple items composed of simple visual features (color, orientation) and abstract shapes (circles, polygons) for brief durations, as these conditions and stimuli are believed to best isolate visual working memory capacity separate from rehearsal, chunking, and passive memory systems (Cowan, 2001). One major conclusion from studies using such stimuli is that the capacity to maintain visual information is strictly limited, often quantified in terms of the number of objects (Luck & Vogel, 1997; Cowan, 2001; Adam, Vogel, & Awh, 2017; Awh & Vogel, 2024), or in terms of a fixed resource pool within each simple visual feature dimension that is distributed among the encoded items (e.g., Bays et al., 2009; Bays, 2014).

However, more recent research has demonstrated that working memory capacity is not fixed, but depends on the kind of information stored: Familiar and meaningful stimuli, such as real-world objects, result in increased neural maintenance activity and better behavioral performance compared to abstract and non-meaningful shapes (Brady et al., 2016; Thibeault et al., 2024; Torres et al., 2024; Asp et al., 2021). One explanation for this benefit is that these stimuli allow participants to extract more relevant visual and semantic features (see Chung et al, 2024a for a review); similar to how maintaining both color and orientation allows the storage of more total information than color alone (Luck & Vogel, 2013; Shin & Ma, 2017).

A recent study significantly expanded on these findings, demonstrating that simple features themselves – colors – are better remembered when they are encoded in a meaningful context (Chung et al., 2023a; Chung et al., 2023b). Specifically, in a series of behavioral experiments, participants showed better color memory performance when these colors were presented as parts of real-world objects (e.g., a blue backpack) compared to unrecognizable scrambled shapes, even though the color-object associations were randomly chosen on every trial, and object identity itself was task-irrelevant. These findings provide a challenge to fixed-capacity models of working memory which assume that how much information can be maintained is strictly limited within a single feature dimension.

What explains this improvement? The observed behavioral effects could be supported by *increases* in active working memory usage; or they could be due to *more efficient* coding of real-world objects compared to abstract stimuli, potentially freeing up resources to better remember the colors; or, the performance benefits could be merely due to differences at retrieval, with no changes during active maintenance. To disambiguate between these possibilities, we here examine both the amplitude and stability of neural activity during the maintenance period of a working memory task. Persistent neural activity during maintenance is the hallmark of working memory functioning, and not present when participants rely on other non-active forms of storage for retaining information (Buschman, Siegel, Roy, & Miller, 2011; Vogel & Machizawa, 2004). One particularly strong marker of active working memory is the contralateral-delay activity (CDA), a sustained negativity of the event-related potential (ERP) indexing how much visual information is actively maintained (Vogel & Machizawa, 2004; for a review, see Luria et al., 2016). The amplitude of the CDA tracks active storage, increasing as more information is actively retained (Vogel et al., 2005; Carlisle et al., 2011; Salahub et al., 2019); it’s more sensitive to visual than verbal load (Predovan et al., 2008); and it remains robust to low-level visual changes such as contrast (Ikkai et al., 2010). We capitalize on this well-established neural marker to examine whether better color memory for real-world objects reflects increased, decreased, or unchanged active maintenance. In addition to the CDA, we examined the stability of the pattern of neural activity over the delay, as an index of the robustness of the memory representation across time. Overall, our results suggest that improved color memory is supported by increased neural engagement during encoding and delay.

### Experiment 1: Assessing Color Working Memory Performance

In Experiment 1, we assessed the behavioral performance of color working memory for intact objects and scrambled objects. Previous studies showed that color working memory is significantly improved when the colors are remembered as parts of meaningful real-world objects (Chung et al., 2023a). Here we aimed to replicate these behavioral effects.

## Materials & Methods

The experiment was approved by the Internal Review Boards at University of California San Diego and Dartmouth College. The sample size, hypotheses, analysis plans, and exclusion criteria for Experiment 1 were pre-registered (https://aspredicted.org/fwhh-h82k.pdf).

### Participants

All participants gave informed consent prior to participating in this experiment. The sample size of the EEG investigation was a priori determined to be twenty-four participants (see Methods below). To achieve greater power for the behavioral effect replication, we doubled the planned sample size in Experiment 1 to be forty-eight participants. In total, fifty-eight participants were recruited from UC San Diego online recruitment platform. Participants’ ages ranged from 18 to 35 years. Following the pre-registered exclusion criteria similar to prior works (e.g., Chung et al., 2023a), we excluded participants’ data if their overall behavioral performance across conditions was lower than d’<0.5 or if more than 10% of their trials were excluded.

Individual trials were excluded if the response time was shorter than 200 ms or longer than 5000 ms. This resulted in ten participants’ data being excluded, resulting in data from forty-eight participants for the final analysis.

### Stimuli

For the intact object condition, real-world object images were sampled from the database of Brady et al. (2013). These objects are generally color-neutral in everyday-life and can be assigned any single arbitrary color. Following Chung et al. (2023a), we rotated the stimuli in hue space along a CIE L*a*b* color wheel, so that any object could occur in any color along the wheel. For the scrambled object condition, these object images were then morphed using the diffeomorphic scrambling technique (Stojanoski & Cusack, 2014), similar to previous work (e.g., Chung et al., 2023a; Chung et al., 2023b; Chung et al., 2024b; Thibeault et al., 2024; Brady & Störmer, 2022). This transformation technique is particularly useful as it can make object images unrecognizable while maintaining their basic perceptual features and visual complexity. None of the object stimuli were repeated throughout the experiment. The exact sizes of the stimuli varied slightly, but they were placed in a white area that extended 135 pixels in height and 135 pixels in width. The colors of stimuli on each trial were randomly selected along the 360-degree color wheel with the constraint that they had to be at least 30 degrees apart from each other, including the foil stimulus at test. This was done so that all colors presented in each trial were discernibly different, and there were no overlapping colors causing interference or confusion at test.

### Experimental Procedure

This was an online experiment where participants completed the study with their own devices. On each trial, participants were presented with an array of three stimuli simultaneously evenly distributed around the central fixation cross for 800 ms. On half of the trials, colored real-world objects were presented and on the remaining half of trials, colored scrambled objects were presented. Afterwards a 800-ms delay period followed, displaying a blank screen with the central fixation cross. At test, one stimulus from the encoding period appeared in two different colors around the center of the screen, and participants were asked to indicate which of the two colors they had encoded initially by clicking the corresponding item. Following previous work (Chung et al. 2023a), one of the stimulus choices appeared in the color that matched the encoded color (target) and the other stimulus appeared in a color 180° away from the target color on the color wheel (foil). The foil colors were also at least 30° away from the other stimulus colors in the encoding array, ensuring no overlap in colors among stimuli on each trial. After the 2-AFC task, participants received feedback in the form of a sound. Note that throughout the experiment, participants were never asked to recall the identities of the stimuli and were only asked about their colors. Thus, the experiment could be done without any regard for the identity of the stimuli. Participants completed 270 trials total (135 trials each condition, randomly intermixed). Prior to the experiment, participants viewed a 10 second video of the task, familiarizing themselves with the task design. See Fig. 1A for an illustration of the trial structure.

**Figure 1.**
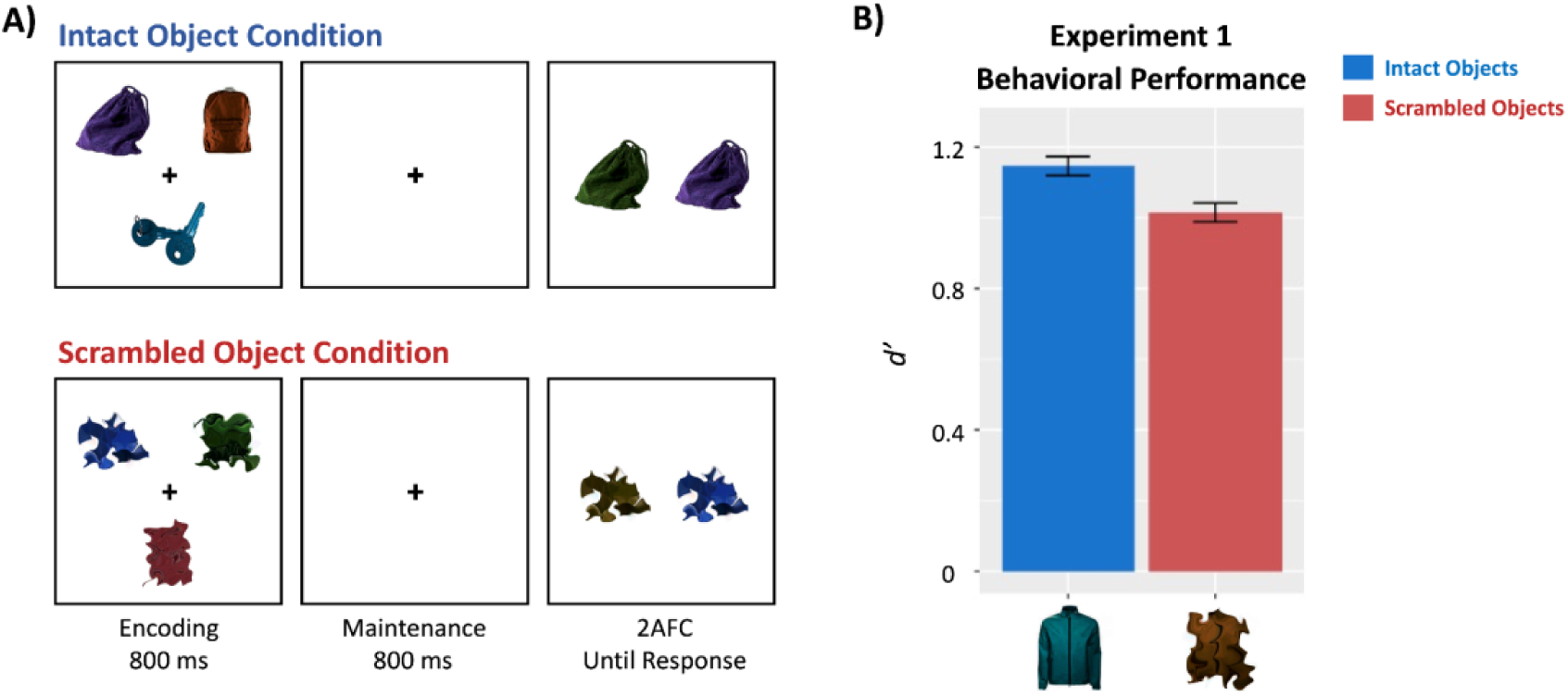
**A)** Trial structure of Experiment 1. All stimulus sizes and distances are for illustration purposes only. **B)** Behavioral performance results of Experiment 1 (N = 48).

### Statistical Analysis

Performance on the color memory task was quantified as d’ for a 2-AFC task (([*z*hits] – [*z*false alarms])/√2) for each participant and condition separately (intact object condition vs. scrambled object condition). The d’ values were then statistically compared using a paired t-test.

## Results

We found a significant increase in color working memory performance for the intact objects (mean d’ = 1.15) compared to scrambled objects (mean d’ = 1.02): *t*(47) = 3.47, p = 0.001, Cohen’s d_z_ = 0.34 (see Fig. 1B). This shows a reliable replication of the previously found meaningfulness benefit in feature working memory (Chung et al., 2023a; Chung et al., 2023b).

### Experiment 2: Assessing Active Working Memory Engagement with EEG

Experiment 1 showed improved color working memory performance for intact real-world objects compared to scrambled objects. In Experiment 2, we used EEG to uncover the active working memory engagement during the maintenance period of the working memory task.

## Materials & Methods

The experiment was approved by the Committee for the Protection of Human Subjects at Dartmouth College.

### Participants

All participants gave written informed consent prior to participating in this experiment. To obtain our goal of twenty-four usable EEG participants, twenty-nine participants were recruited from Dartmouth College using their online recruitment platform. Participants’ ages ranged from 18 to 35 years, and all participants had normal or corrected-to-normal vision. Prior to the experiment participants were given the Ishihara color blindness test (Clark, 1924). Similar to Experiment 1, participants’ behavioral and EEG data were excluded if their overall behavioral performance was lower than d’< 0.5. Their EEG data were excluded if more than 40% of trials were excluded due to EEG artifact rejection. No participant was excluded due to behavioral performance. Five participants were excluded from the EEG analysis due to artifact rejection, totaling our target of twenty-four participants. On average, about 17.7% of trials were rejected due to EEG artifacts across the remaining participants. The final sample size of 24 for the EEG analysis was determined following protocols of previous studies investigating CDA measures of meaningful stimuli (e.g., 19 participants Asp et al., 2021; 18 participants in Brady et al., 2016; 22 participants in Thibeault et al., 2024).

### Stimuli

Stimuli used in Experiment 2 were similar to Experiment 1, except all 540 real-world object images were sampled from Brady et al. (2013). On a given trial, three colored objects appeared on each side of the screen. Two of the stimuli were centered 4.1° of visual angle away from the fixation along the horizontal meridian and +/-3.5° of visual angle along the vertical meridian, while the other one was centered 9.7° away from fixation along the horizontal meridian, centered on the vertical meridian (see Fig 2A for an illustration). The exact stimulus size varied slightly across objects, but they were all centrally placed in a white area that extended 5.2° in height and 5.2° in width in visual angle for the two of objects at the top and bottom locations, and 5.7° in height and 5.7° in width for the object presented along the horizontal meridian that was more distant from fixation. Similar to Experiment 1, the colors of objects on each side of the screen were randomly selected along the 360-degree color wheel with the constraint that they had to be at least 30 degrees apart from each other. Due to the limited number of objects sampled, some objects repeated throughout the experiments. However, none of the objects repeated more than once, and this was the same across the two conditions.

**Figure 2.**
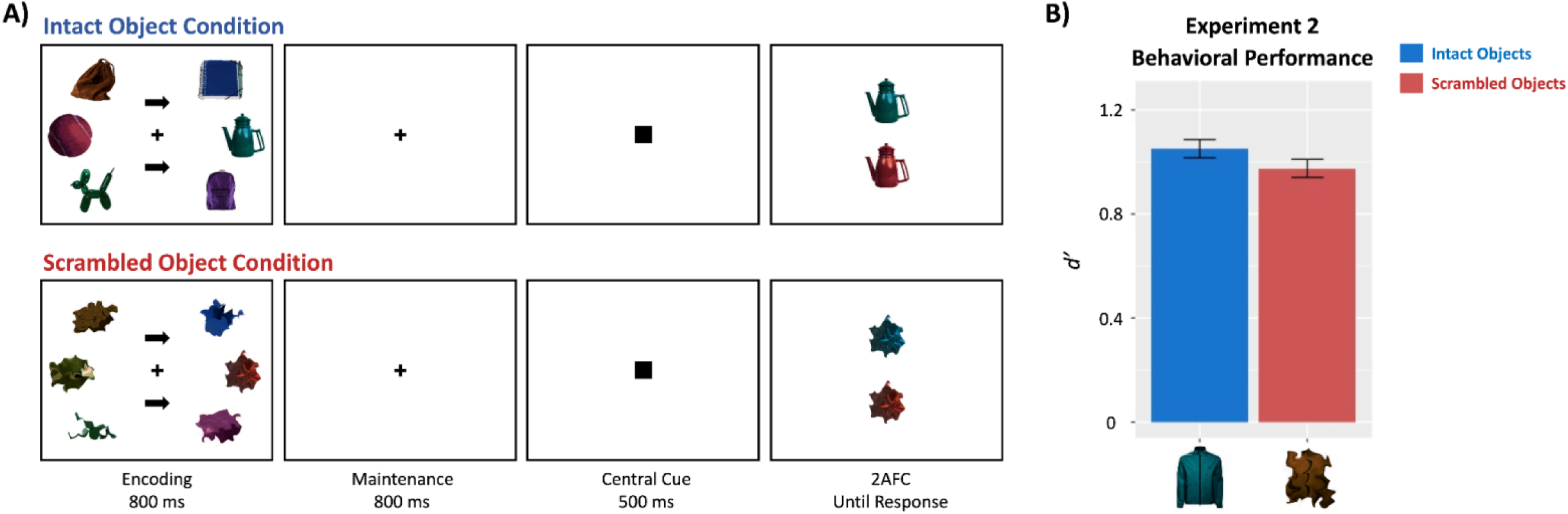
**A)** Trial structure of Experiment 2. All stimulus sizes and distances are for illustration purposes only. **B)** Behavioral performance results of Experiment 2 based on the 24 participants used in the main EEG analysis.

### Experimental Procedure

Similar to Experiment 1, participants were instructed to remember the colors of three stimuli over a brief delay period. Throughout each trial, a small black fixation cross was presented in the center of a white screen and participants were instructed to maintain fixation. We used a lateralized memory display to allow the measurement of the lateralized CDA component. Thus, the memory encoding display consisted of three stimuli presented on each the left and right sides of the screen along with two black arrow cues at the center of the screen (one above and one below the fixation cross) pointing towards the to-be-remembered side (left or right). On half of the trials, colored real-world objects were presented and on the remaining half of trials, colored scrambled objects were presented. Participants were instructed to remember the colors of the stimuli on the side that the arrows pointed to while maintaining their gaze on the central fixation cross. After a 800-ms encoding period, a 800-ms delay period followed displaying a blank screen with just the central fixation cross, followed by a probe (a small black square, 0.9° height and width; 500ms) that appeared at the center of the screen, prompting participants of the beginning of the response period. Subsequently, participants performed the 2-AFC task where one stimulus from the to-be-remembered side appeared in two different colors around the center of the screen (slightly shifted along the vertical meridian), and participants were asked to indicate which of the two colors they had encoded initially by pressing up- and down-arrow keys on the keyboard corresponding to the relative position of the items on the screen (top vs. bottom). The stimuli for the 2-AFC task were chosen the same way as in Experiment 1. After the response, participants received feedback in the form of a fixation cross color change (green correct, red incorrect) after which the screen turned fully white. After each trial, participants could start the next trial with a button press. The next trial then began after an inter-trial interval (ITI) of 800ms (first 19 participants) or randomly jittered 600-1,000ms (last 5 participants). Participants completed 432 trials in total (216 trials each condition, randomly intermixed), and also went through a practice block of 10 trials prior to the main experiment. See Fig. 2A for an illustration of the trial structure.

### Electrophysiological recording

EEG was recorded continuously from 32 Ag/AgCl electrodes (arranged in the 10-20 system) mounted in an elastic cap and amplified by an ActiCHamp amplifier (BrainProducts, GmbH). The horizontal electrooculogram (HEOG) was recorded from two additional electrodes positioned on the external ocular canthi which were grounded with another electrode placed on the neck of the participant. All scalp electrodes were referenced to the right mastoid online and were digitized at 500 Hz. EEG data was filtered with a bandpass of 0.01–112.5 Hz online.

### Statistical Analysis

#### Behavioral data analysis

The behavioral analysis was identical to Experiment 1.

#### EEG Analysis

All EEG data analyses were performed using EEGLAB (Delorme & Makeig, 2004) and ERPLAB (Lopez-Calderon & Luck, 2014) toolboxes and custom-written scripts. Data were epoched into trials aligned to the onset of the encoding display. Artifacts were detected using an automated pipeline and artifact rejection was performed from 200 ms before the onset of memory stimuli to 1600 ms afterwards (the end of the delay period). Trials with eye blinks (peak-to-peak at channel FP1, threshold at 150 μV), excessive eye movements (step function at HEOG, threshold at 17 μV which corresponds to ∼1° of saccade; cf., Luck, 2014; Hillyard & Galambos, 1970), and large noise (peak-to-peak all channels, threshold at 300 μV) were excluded from the analysis. The average rejection rate of participants included in the final analysis was 17.7%. Artifact-free data were re-referenced to the average of the left and right mastoids, digitally low-pass filtered at 30 Hz (Butterworth filter, 12dB/octave roll-off), and baseline corrected to the 200 ms prestimulus interval.

#### Parietal-occipital lateralized ERPs

We examined lateralized activity over posterior electrode sites both during the encoding (400-800 ms) and delay (1200-1600 ms) periods. Our main interest was to assess whether remembered colors of real-world objects vs. scrambled objects result in a difference in the delay period (the CDA) as this marks the active maintenance of visual representations (Vogel & Machizawa, 2004; McCollough et al., 2007; Unsworth et al., 2015; Adam et al., 2018). However, based on recent findings indicating that lateralized differences may emerge during the encoding period when using relatively long encoding times and meaningful stimuli (cf., Asp et al., 2021; Thibeault et al., 2024), we included both time intervals in our main analyses. EEG epochs were averaged separately for each object condition (intact vs. scrambled) and hemifield (remember left vs. right) and were then collapsed across to-be-remembered visual hemifield (left/right) and hemisphere of recording (left/right) to obtain waveforms recorded over the hemisphere contralateral and ipsilateral with respect to the to-be-remembered side. To quantify activity that was related to the encoding and memorization of the cued items in particular, the ipsilateral waveform was subtracted from the contralateral waveform. Thus, this difference wave reflects the increase in neural activity that’s specific to the to-be-remembered items, removing all purely sensory-related activity. The mean amplitude for each participant was calculated by averaging activity across four posterior-occipital electrode pairs (PO3/PO4, PO7/PO8, P7/P8, and O1/O2; Asp et al., 2021) from 400 to 800 ms post stimulus onset for the encoding period and 400 ms to 800 ms after the stimuli offset (1200-1600 ms post encoding display onset). Electrode sites and CDA time windows were chosen a-priori based on previous studies (see Roy & Faubert, 2023 for a review); the encoding period time window was chosen to avoid early visually-evoked potentials (i.e., P1/N1) and match the duration of the CDA analysis. For each time window, mean amplitudes were compared across conditions (intact vs. scrambled) using paired t-tests.

We also examined differences in the topographies across the two conditions during both encoding and delay time periods. Scalp topographies of the lateralized encoding and retention periods are plotted in Figure 3D as the contralateral-minus-ipsilateral difference waveform, projected on the right side of the scalp. To statistically compare these topographies, we first normalized the difference waveform by dividing the amplitude from each electrode site by the square root of the sum of the squared voltages across all electrodes within each condition (McCarthy & Wood, 1985) to ensure differences in overall amplitudes would not drive the differences in results. Then we ran a 2X2 ANOVA with 13 lateralized electrode pairs and two conditions (intact vs. scrambled objects) as within-subjects factors.

**Figure 3.**
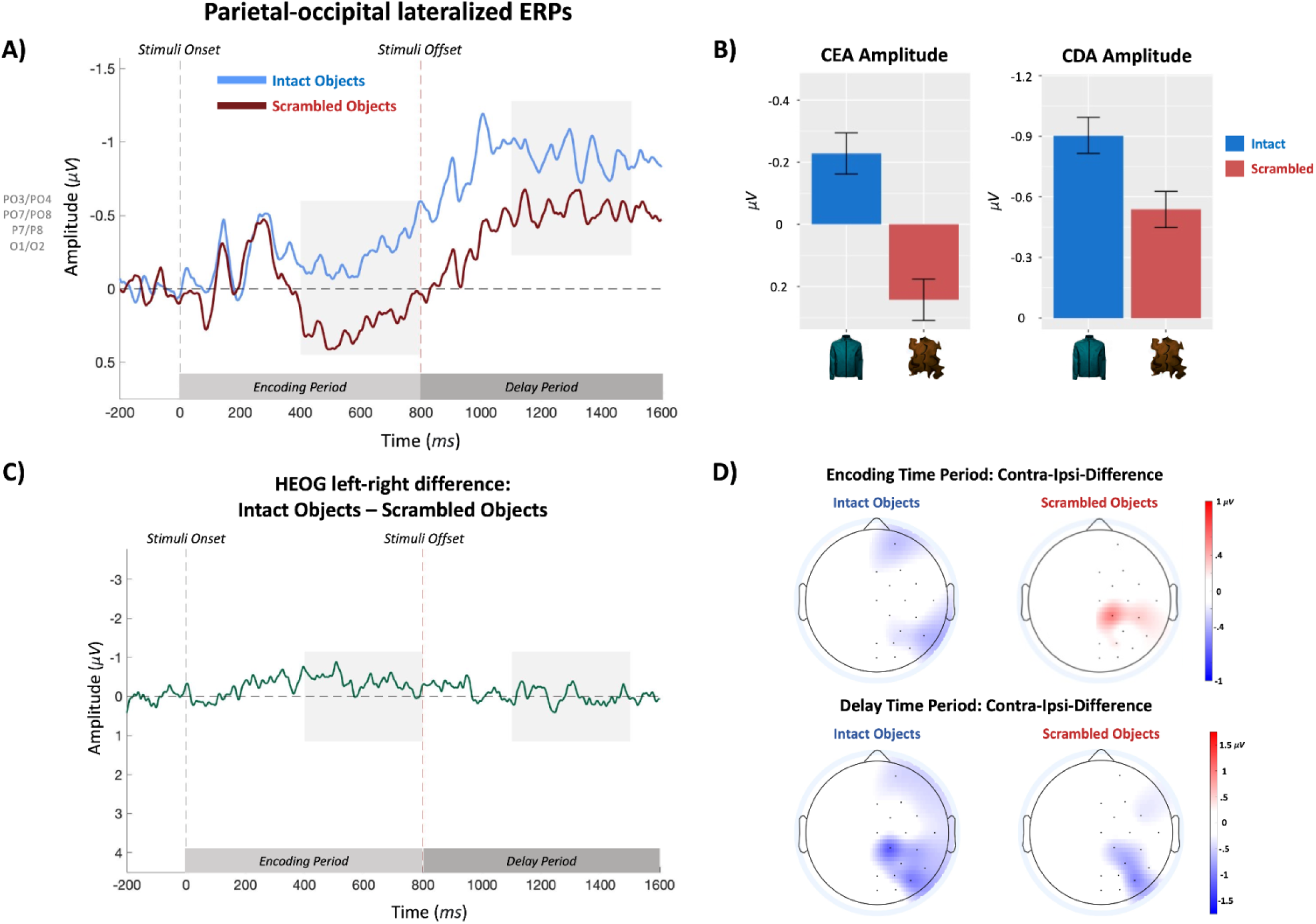
**A)** Contralateral-minus-ipsilateral waveforms averaged over parietal-occipital electrodes separately for intact (light blue) and scrambled (dark red) object conditions. Highlighted areas depict the time windows for the ERP analyses for the contralateral-encoding (CEA) and contralateral-delay activities (CDA). **B)** Mean CEA and CDA amplitudes separately for each condition. Error bars reflect within-subject standard errors of the mean. **C)** HEOG subtraction waveforms. Highlighted areas are the time windows for CEA and CDA analyses. **D)** Spatial topographies of the contralateral - minus - ipsilateral waveforms during the encoding and delay time periods. Mean difference is projected on the right side of the scalp. Note that the scale differs between encoding (top) and delay (bottom) time periods.

#### Cross-Temporal Pattern Similarity

To evaluate the neural stability of the memory representations for each condition, we adopted a novel analysis method (based on Spaak et al., 2017; see also Liu et al., 2020) comparing the pattern similarity of the lateralized difference waveforms across time. Specifically, separately for each condition, we computed the cosine similarities of the activity pattern of these ERP waveforms over time, smoothed over 10ms. We separated this similarity analysis into two sets of electrodes, one using a set of parietal-occipital electrodes (P7/P8, P3/P4, PO3/PO4, PO7/PO8, O1/O2), and one using a set of fronto-central electrode (FP1/FP2, F3/F4, FC1/FC2, FC5/FC6, C3/C4); given that this was a *visual* working memory task, we expected the posterior set of electrodes to be more sensitive to the condition differences. Figure 4A depicts the resulting cross-temporal analysis matrices which show the neural similarity time by time across the entire trial period, separately for posterior (top) and anterior (bottom) scalp sites. Higher cross-temporal similarity reflects more stable representations. Across all conditions, we observed an increase in similarity across time. The critical question was whether this increase in neural stability arose at different points in time for the intact vs. scrambled object condition. To test this, we collapsed the cross-temporal similarity matrices by averaging the similarity values between a given time point and the average of all later time points (i.e., mean cosine similarities of *t*_1_ vs. *t*_2_-*t*_n_; *t*_2_ vs. *t*_3_-*t*_n_, etc.). We then statistically compared the time courses of the average similarity values across conditions using a cluster-based permutation t-test. At each time bin (10ms), we performed a paired t-test comparing the similarity values for the real and scrambled conditions across participants. Contiguous time bins with uncorrected p-values below 0.05 were grouped into clusters. For each cluster, we summed the t-values to create a cluster-level test statistic. To assess significance, we generated a null distribution by randomly permuting the time bin labels within each subject 1000 times and recomputing the t-tests and cluster-level statistics for each permutation. The p-value for each observed cluster was computed as the proportion of permuted cluster-level statistics that exceeded the observed cluster sum. This procedure controls for multiple comparisons while preserving the temporal structure of the data.

**Figure 4.**
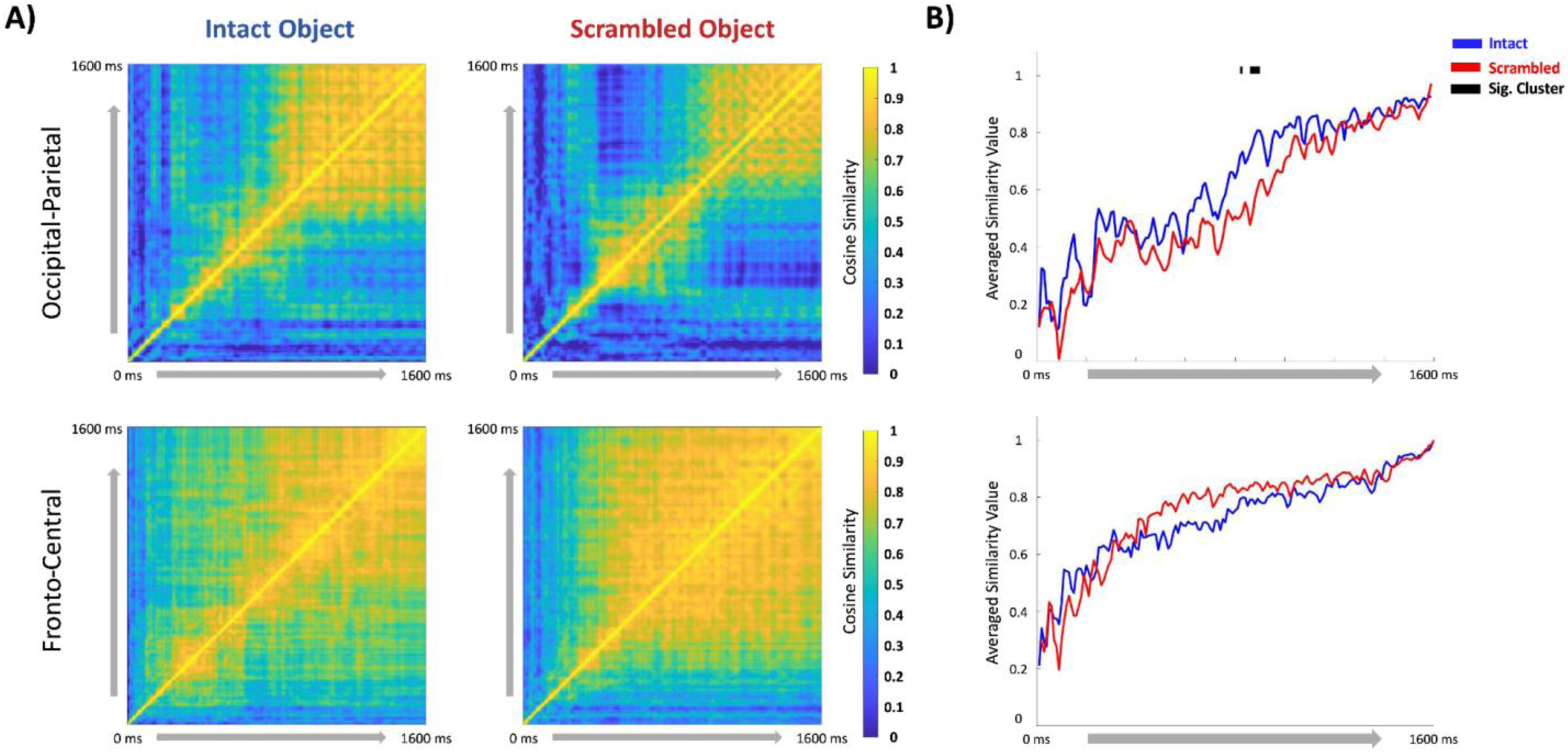
**A)** Cross-temporal stability matrices depicting cosine similarities of lateralized ERP waveform patterns across time. Matrices on top are parietal-occipital electrode sites while the bottom ones are fronto-central electrode sites. **B)** Cosine similarity values between each time window and all later time windows were averaged to compare the time course of stability across the two conditions. Generally, ERP patterns became more stabilized over time, indicated by upwards trends of the averaged similarity values. For parietal-occipital sites, the neural activity pattern for intact objects became stable earlier compared to scrambled objects, as indicated by the significant difference around 800-900ms.

#### Bilateral fronto-central bilateral ERPs

As an exploratory analysis, we also compared non-lateralized neural activity between the two experimental conditions at fronto-central electrode sites (FC1, FC2, CP1, CP2, Cz) during both the encoding and delay periods using the same time windows as for the lateralized ERPs (400-800 ms and 1200-1600 ms post stimulus onset). The mean ERP amplitudes between the two conditions were compared using a paired t-test. Note that such global differences in bilateral activity is controlled for in the main CDA analysis as non-lateralized activity is subtracted out.

## Results

### Behavioral Results

We first analyzed the behavioral data including all the participants who met the behavioral inclusion criteria (all 29 participants). This revealed the expected increase in color working memory performance for the intact objects (mean d’ = 1.09) compared to scrambled objects (mean d’ = 0.98): *t*(28) = 2.25, p = 0.03, Cohen’s d_z_ = 0.43, replicating Experiment 1 and other previous work (Chung et al., 2023a; Chung et al., 2023b). When only including the 24 participants that were retained for the EEG analysis after artifact rejection, this effect remained numerically present though was statistically non-significant (mean d’ = 1.05 for intact objects vs. 0.98 for scrambled *t*(23) = 1.54, p = 0.14, Cohen’s d_z_ = 0.32 (see Fig. 2B).

### Contralateral Encoding Activity (CEA)

All EEG analyses focused on the 24 participants who met the EEG inclusion criteria. During the encoding period (400-800ms), colors superimposed on intact objects elicited a larger contralateral vs. ipsilateral negativity (mean amplitude = −0.23 μV) relative to colors superimposed on scrambled stimuli (mean amplitude = 0.24 μV): *t*(23) = 5.03, p < 0.001, Cohen’s d_z_ = 0.89 (see Fig. 3A/3B).

Statistical analysis of the spatial topography during encoding resulted in a significant interaction between the memory conditions and electrode sites (*F*(12, 276) = 2.42, p = 0.005, η_p_ ^2^ = 0.095), indicating a shift in the pattern of activity across conditions. There was also a significant main effect of condition (*F*(1, 23) = 5.18, p = 0.03, η_p_ ^2^ = 0.18) but no main effect of electrode locations (*F*(12, 276) = 1.42, p = 0.15, η_p_ ^2^ = 0.06). Visual inspection suggests that the intact object condition showed a more lateralized parietal-occipital focus, whereas the scrambled object condition showed a more central-parietal focus of activity (see Fig. 3D).

### Contralateral Delay Activity (CDA)

During the retention period, mean CDA amplitude (1200-1600 ms) was significantly more negative (mean amplitude = −0.90 μV) for the intact object condition than the scrambled object condition (mean amplitude = −0.54 μV): *t*(23) = 2.90, p = 0.008, Cohen’s d_z_ = 0.59 (see Fig. 3A/3B), indicating that active maintenance activity was increased for colors superimposed on real objects at encoding. This pattern was present when including all 29 participants’ data (see Supplemental Material).

The topographical analysis revealed a significant interaction between the memory conditions and electrode sites (*F*(12, 276) = 2.003, p = 0.02, η_p_ ^2^ = 0.08), indicating a reliable difference in the spatial distribution of activity, similar as during the encoding period. There was no main effect of condition (*F*(1, 23) = 1.23, p = 0.28, η_p_ ^2^ = 0.05), but a significant main effect of electrode location (*F*(12, 276) = 3.89, p < 0.001, η_p_^2^ = 0.15). Visual inspection of the spatial topographies suggests that the activity pattern was more spread-out towards parietal-central and temporal electrode sites for the intact object condition compared to the scrambled condition, in addition to increasing in amplitude (see Fig. 3D).

### Cross-Temporal Pattern Similarity

The cross-temporal matrices for intact vs. scrambled object conditions, separately for parietal-occipital and fronto-central electrode channels, are shown in Figure 4A. Overall, as time evolves, neural stability increases across both conditions. Visual examination of these cross-temporal matrices suggests that the neural pattern became stable earlier for intact vs. scrambled objects, especially over parietal-occipital electrode sites. Our statistical analysis examining the time course of neural stability supported this, resulting in a cluster of significant differences in stability between the two conditions around 800∼900 ms for the occipital-parietal electrode sites (see Fig 4B, top). No reliable differences in neural stability between the two conditions were found for the fronto-central electrode sites (Fig 4B, bottom).

### HEOG

To ensure that the observed lateralized effects (e.g., CEA, CDA) were not attributable to systematic eye movement differences between conditions, we analyzed the HEOG waveforms for each cued side (left/right) across the object conditions (intact/scramble) during both the CEA and CDA time windows. Specifically, we subtracted the HEOG activity of the remember-right condition from the activity of the remember-left condition, separately for each object condition; then, we subtracted the scrambled object condition from the intact object condition. These subtracted HEOG waveforms were then tested against zero using single-sample t-tests. The results indicated that the subtracted HEOG waveforms did not significantly deviate from zero in either time window: (CEA: *t*(23) = 1.07, p = 0.29; CDA: *t*(23) = 0.02, p = 0.98; see Fig. 3C).

This suggests that the observed differences in CEA and CDA across the conditions were not driven by differences in systemic eye movements.

### Fronto-Central Negativity during Encoding and Delay Period

We observed a robust non-lateralized ERP difference over fronto-central electrode sites across the intact and scrambled object conditions. Specifically, the intact object condition elicited a large negative slow-wave relative to the scrambled object condition during the encoding period (*t*(23) = 9.25, p < 0.001, Cohen’s d_z_ = 0.69), which sustained throughout the delay period (*t*(23) = 4.06, p < 0.001, Cohen’s d_z_ = 0.32; see Fig. 5A/5B/5C). This suggests that activity over fronto-central electrode sites tracks the meaningfulness of items at encoding and during the delay.

**Figure 5.**
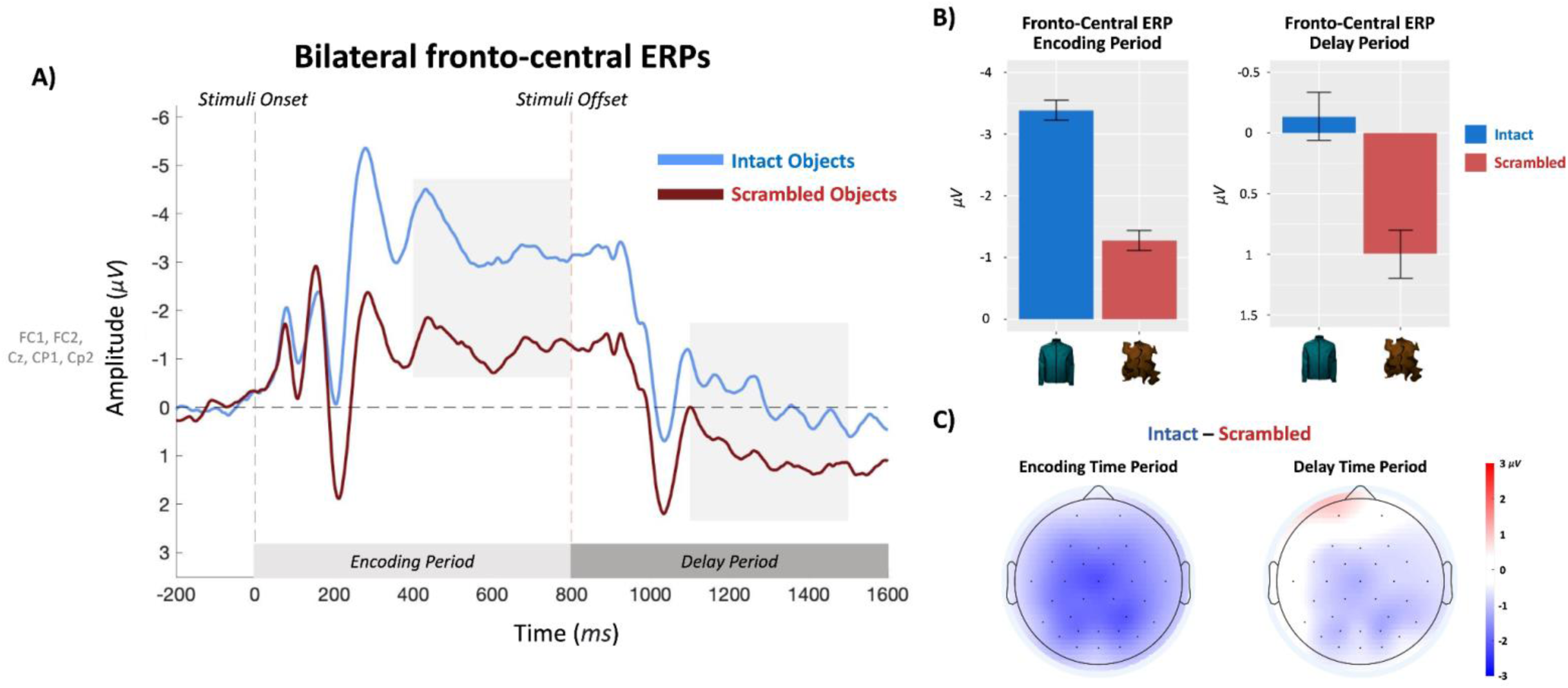
**A)** Non-lateralized waveforms recorded over fronto-central electrode sites separately for intact and scrambled objects. Highlighted areas indicate the time windows for statistical analyses. **B)** Mean ERP amplitudes during the encoding and delay time periods. **C)** Spatial topographies of the intact - minus - scrambled object conditions for each time window.

## General Discussion

We recently found improved working memory performance for colors when they were encoded as parts of meaningful objects, relative to colors that were superimposed on non-meaningful shapes that were matched in visual complexity (Chung et al., 2023a, Chung et al., 2023b; see also Loaiza et al., 2024). Here, we demonstrate that this performance benefit is accompanied by increased neural delay activity and earlier-arising neural stability for intact objects compared to scrambled objects. This finding of increased neural engagement contrasts with alternative theories, like efficient coding of color information (which would result in reduced memory engagement), or non-active processes such as more effective retrieval at test from passive memory systems, like activated long-term memory (e.g., Cowan, 1988).

We also observed differences in the spatial topography across conditions: the CDA appeared to be more spread-out throughout parietal, central, and temporal sites in the intact object compared to the scrambled object condition that showed a more occipital focus. Such shifts in CDA topography may indicate recruitment of more task-relevant neural regions in the intact object condition during the encoding and maintenance periods (i.e., sensory recruitment; Adam et al., 2022; for similar shifts in CDA from working memory task relevance, see Breitinger et al., 2022). For example, it has been shown that colors of real-world objects activate not only low-level color maps (e.g., V4), but also areas in higher-level visual regions that are tuned to objects and their associated colors (e.g., Bramão et al., 2010; Lafer-Sousa et al., 2016). Thus, simple features like colors may be coded – and maintained – redundantly across the visual hierarchy, creating stronger, more stable representations. Further, associating simple features like color to real-world objects may activate additional semantic knowledge structures of these objects, effectively increasing memory dimensionality and thereby reducing interference (Wyble, 2016). This interpretation is consistent with other recent studies showing increased neural maintenance activity and higher performance when participants remember not just simple visual features but entire identities of real-world objects (Brady et al., 2016; Thibeault et al., 2024; Asp et al., 2021). The current results go beyond these previous findings by demonstrating that increases in representational dimensionality due to links to semantic knowledge not only improve memory for the objects themselves but also provide an effective scaffold for memorizing their low-level and abstract visual features (Chung et al., 2023a; b). Overall, this suggests that expanded neural recruitment enables higher-dimensional working memory representations that support the maintenance of even simple features, either through redundant coding and/or by connecting lower-level visual features to semantic meaning in a hierarchically structured way.

Is it possible that the observed increase in CDA reflects memory for the intact objects themselves, and not their colors? Such an account would require that this study be unique in finding both a behavioral benefit for a condition and a CDA benefit for the same condition, but those benefits somehow being due to different causes — in direct conflict with repeated findings of a strong relationship between CDA and behavior at both the group and individual level (Vogel & Machizawa, 2004; Störmer et al., 2013, Roy & Faubert, 2022; Luria et al., 2016). This seems unlikely, especially given several previous studies demonstrating that the CDA component represents storage of task-relevant information only (Woodman & Vogel, 2008; Zhu et al., 2022). For instance, Zhu et al. (2022) showed that features that are used for the selection in working memory tasks but are not tested at retrieval are not reflected in the CDA. Similarly, Williams & Drew (2020) reported that processing of task-irrelevant real-world objects was not reflected in the CDA (Exp. 1, Williams & Drew, 2020). Thus, while it seems likely that some aspects of object information beyond color are incidentally represented in memory (Chung et al., 2024b), the current results seem most compatible with the interpretation that the observed CDA increase is not merely due to active maintenance of task-irrelevant object-identity features, but reflects a boost in maintenance capacity for the task-relevant colors — consistent with behavioral findings that people are better at remembering colors presented on real objects even if those objects are not presented at test (Exp. 5 in Chung et al., 2023a).

The present study used carefully generated control stimuli that matched the low-level visual features and complexity of real-world objects, to ensure that observed neural differences cannot be attributed to differences in visual complexity. This is particularly important as some previous studies have shown CDA amplitudes may be sensitive to large perceptual differences among stimuli (e.g., Woodman & Vogel, 2008; Rajsic et al., 2018). However, more complex but meaningless stimuli actually resulted in lower CDA amplitudes than simple single features, consistent with lower behavioral performance for such stimuli (Luria et al., 2010; Gao et al., 2009); only stimuli that are both complex and meaningful resulted in higher CDA amplitudes compared to abstract stimuli, again consistent with higher behavioral performance for such stimuli (Asp et al., 2021; Thibeault et al., 2024; Brady et al., 2016). This suggests that the CDA may reflect a broader neural marker of general memory strength in visual working memory (i.e., indexing how much information is being held in visual working memory more generally; Salahub et al., 2019; Gao et al., 2009), and is not just an assessment tool for discrete item limits (Ikkai et al., 2010; Fukuda et al., 2010; Balaban et al., 2019; Hakim et al., 2019).

We also observed earlier stabilization of neural activity patterns in the real-world object condition. This may reflect shifts in how working memory representations form over time. For instance, stable representation may emerge more rapidly for meaningful objects due to readily available semantic structures that facilitate mapping the memoranda onto those higher-level structures. In contrast, meaningless objects may require longer time to reach a stable neural representation, resulting in greater pattern shifts. While further work is needed to interpret this novel approach to analyzing lateralized ERPs in working memory (i.e., testing different set sizes or chunking effects), our findings open a new direction for CDA research.

We found that the contralateral negativity emerged earlier for real-world objects, with a larger effect already present during the encoding period, before the CDA window. This matches other recent studies that also reported lateralized ERP differences emerging during encoding when comparing working memory for meaningful vs. non-meaningful stimuli (Thibeault et al., 2024; Asp et al., 2021). This could reflect faster and more accurate encoding of the stimuli into working memory, as well as limits in how much information can be extracted and maintained during perception (Tsubomi et al., 2013; Balaban & Luria, 2019; for a review see Emrich, 2022). Such early condition differences contrast with earlier claims in the literature of no effects of encoding time (Luck & Vogel, 1997; Alvarez & Cavanagh, 2004), but are consistent with newer work showing repeatedly that encoding format and time can play a critical role in working memory processes for simple stimuli (Schurgin et al., 2020; Quirk et al., 2020) and meaningful objects (Brady et al., 2016; Brady & Störmer, 2023; Chung et al, 2023b). The present data provide further support that memory capacity depends on limits that can be observed even when the stimuli are still on the screen – and that real-world objects can provide a useful structure to initially build, and then maintain, robust representations.

Finally, our study also reveals a novel neural signature that appears to track processes related to recognizing an intact vs. scrambled object. Beginning during encoding and continuing through delay, we observed a large non-lateralized slow-wave over fronto-central electrode sites that was more negative for the intact vs. scrambled condition. This sustained negativity for recognizable vs. completely scrambled objects resembles an ERP finding in the word reading literature: the N400, a well-known marker related to semantic understanding (Kutas & Hillyard, 1980), is more negative for pseudowords compared to words (as well as semantically inconsistent vs. consistent words), but interestingly, has also been reported to be more *positive* for fully scrambled nonwords (Bentin et al., 1999; Ziegler et al., 1997; Coch & Mitra, 2010).

This is consistent with our results showing that fully scrambled objects result in a relative positivity compared to intact objects. This suggests that amplitude changes in the N400 component, traditionally linked to word processing, may reflect broader cognitive processes that distinguish between semantically meaningful versus semantically inconsistent and, critically, also fully scrambled stimuli across domains (verbal and visual). While further research is necessary to fully understand what cognitive processes underlie the observed amplitude differences between intact vs. scrambled objects over fronto-central sites, they appear to robustly distinguish between these stimulus conditions. ^1^

In conclusion, our results show that working memory for simple features is shaped by the context these features are encoded in. Meaningful context at encoding – being part of a real-world object – promotes stronger neural engagement and earlier temporal stability of color memory representations during encoding and maintenance processes. Broadly, this indicates that working memory processes, and consequently its capacity, are flexible, even with respect to a single feature dimension.

## Acknowledgements

Special thanks to Sadye Law, our research assistant who helped with initial data collection and pilot experiments, and Alison Sasaki and Sam (Seho) Jung who helped finish data collection. TFB is supported by the National Science Foundation grant 2141189.

## Supplemental Material

### EEG Analyses including all 29 participants

For completeness, we have run the same main analysis for CDA amplitudes including all 29 participants. The 5 participants who were excluded for main analyses had on average 54.12% of trials excluded due to EEG artifacts. Same pattern of results emerged with all 29 participants: the CDA amplitude was larger for the intact object condition than the scrambled object condition (t(28) = 2.64, d_z_ = 0.45, p = 0.01).

## Footnotes

1 A similar pattern of non-lateralized ERP result was observed in a previous study investigating visual working memory of real-world objects and scrambled objects (Thibeault et al., 2024; personal correspondence).

